# Novel features of miRNA and isomiR-mRNA interactions

**DOI:** 10.1101/2025.05.12.653582

**Authors:** Matthew Weston, Rony Chowdhury Ripan, Xiaoman Li, Haiyan Hu

## Abstract

Studying the interactions between microRNAs/isomiRs and mRNAs is critical due to their fundamental roles in gene regulation and their involvement in various diseases. Although many isomiRs have been identified, the analysis of their interactions with mRNAs remains in its early stages. In this study, we compiled available human chimeric read data, each comprising a microRNA or isomiR segment paired with an mRNA fragment and identified 1,747 isomiRs and over 5 million microRNA/isomiR–mRNA interactions. We observed that microRNAs with higher adenine and thymine content, and lower cytosine content, tend to have more isomiRs and more mRNA targets. Notably, an average of 18.9% of mRNA targets were bound exclusively by isomiRs, not by their microRNA counterparts. Furthermore, isomiRs sharing the same seed sequences as their reference microRNAs may bind to different targets from their microRNAs, highlighting functional divergence. Interestingly, 20.0% of microRNAs and 8.2% of isomiRs appear to bind mRNAs independently of their seed regions. Among those that do utilize seed regions, 94.5% of microRNAs and 95.7% of isomiRs also engage non-seed regions, suggesting a broader and more complex binding behavior. Our findings offer new insights into microRNA/isomiR–mRNA interactions.

## Introduction

IsomiRs are isoforms of their reference microRNAs (miRNAs). They arise from alternative miRNA biogenesis processes involving nucleotide additions, deletions, or substitutions, often resulting in differences in length and sequence compared to their miRNA counterparts^1^. Like miRNAs, isomiRs regulate gene expression by binding to mRNA targets, leading to mRNA degradation or translational repression^2^. However, due to changes in their seed regions, isomiRs are shown to have distinct mRNA targets from their reference miRNAs, thereby altering gene expression programs in normal physiological conditions and disease states^3^. It is thus important to study isomiRs, especially to understand their targeting.

Previous studies have investigated the altered mRNA targets by isomiRs compared with their reference counterparts^2-4^. They have also explored their potential as biomarkers for disease detection, prognosis, and diagnosis^5-8^. In addition, several databases have been developed to store and organize isomiR-related data, such as isomiRdb, IsomiR Bank and the tumor isomiR encyclopedia (TIE)^9-11^. For instance, isomiRdb has cataloged over 70,000 isomiRs across approximately 43,000 samples^9^.

Despite these efforts, to our knowledge, only a few studies have systematically examined isomiR-mRNA target interactions. One such effort is isomiRTar^12^, which analyzed the predicted targets of 1,022 5’ isomiRs across 31 cancer types using tools such as miRDB^13^ and targetScan^14^. These tools predict mRNA targets based on sequence features and negative co-expression patterns of isomiRs and their targets. However, these tools were originally developed for miRNAs and may not be optimal for isomiR targeting. Another effort is DMISO^15^, a deep learning model trained on crosslinking, ligation and sequencing of hybrids (CLASH) data^16^. CLASH captures direct miRNA/isomiR–mRNA interactions by generating chimeric reads composed of physically ligated miRNA or isomiR fragments and their mRNA target segments, providing context-specific interaction evidence. While DMISO represents a step forward, it was trained on only six CLASH samples available at the time, limiting its generalizability. Improving prediction models requires both more data and a deeper understanding of miRNA/isomiR binding mechanisms.

In the past decade, additional CLASH-like technologies have emerged, including CLEAR-CLIP, Quick CLASH (qCLASH), and chimeric e-CLIP, which produce high-resolution chimeric reads of miRNA/isomiR-mRNA interactions^17-19^. These additional data provide an unprecedented opportunity to systematically investigate the dynamics and specificity of miRNA/isomiR targeting across conditions and cell types.

In this study, we analyzed 1,747 isomiRs and 5,556,801 miRNA/isomiR-mRNA interactions derived from 67 publicly available CLASH-like human samples across six studies. Of the isomiRs identified, 73.1% were supported by curated entries in isomiRdb^9^, while the remaining 26.9% likely represent novel isomiRs, exhibiting similar frequencies of occurrence in the analyzed samples. Moreover, approximately 6.7% of these miRNA/isomiR-mRNA interactions were previously recorded in miRTarBase^20^. Notably, we found that miRNAs with more isomiRs and mRNA targets tend to have higher adenine and thymine content and lower cytosine content. These findings highlight novel sequence characteristics associated with broader target engagement of miRNAs/isomiRs.

## Material and Methods

### Process CLASH and CLASH-like data

We obtained a total of 67 human CLASH and CLASH-like samples across six datasets, which were the only available samples we could access and process for chimeric reads. These data included nine samples from CLASH^16^, twelve from CLEAR-CLIP^19^, nine from qCLASH^17^, thirteen from chimeric eCLIP^18^, three from Kozer et al.^21^, and twenty-one from Fields et al.^22^.

For each sample, we followed the procedure outlined in DMISO^15^ to process the reads and extract miRNA/isomiR-mRNA interactions, closely mirroring the methodology used in the original CLASH study^16^ (Fig. 1). Briefly, we began by removing adaptors from the reads and filtering low-quality reads with trimmomatic^23^ and fastqc. We then eliminated duplicate reads using the ShortRead^12^ library in R. Next, we mapped the processed reads to the protein-coding transcript sequences from GENCODE^24^ (version 46) and the human mature miRNA sequences from miRBase^25^ (version 22.1) using BLAST^26^ (version 2.15.0+). Mapping was performed with an E-value threshold of 0.1 and a default word size of 11. Reads aligning to the antisense strand or containing alignment gaps were excluded from further analysis.

**Fig 1.**
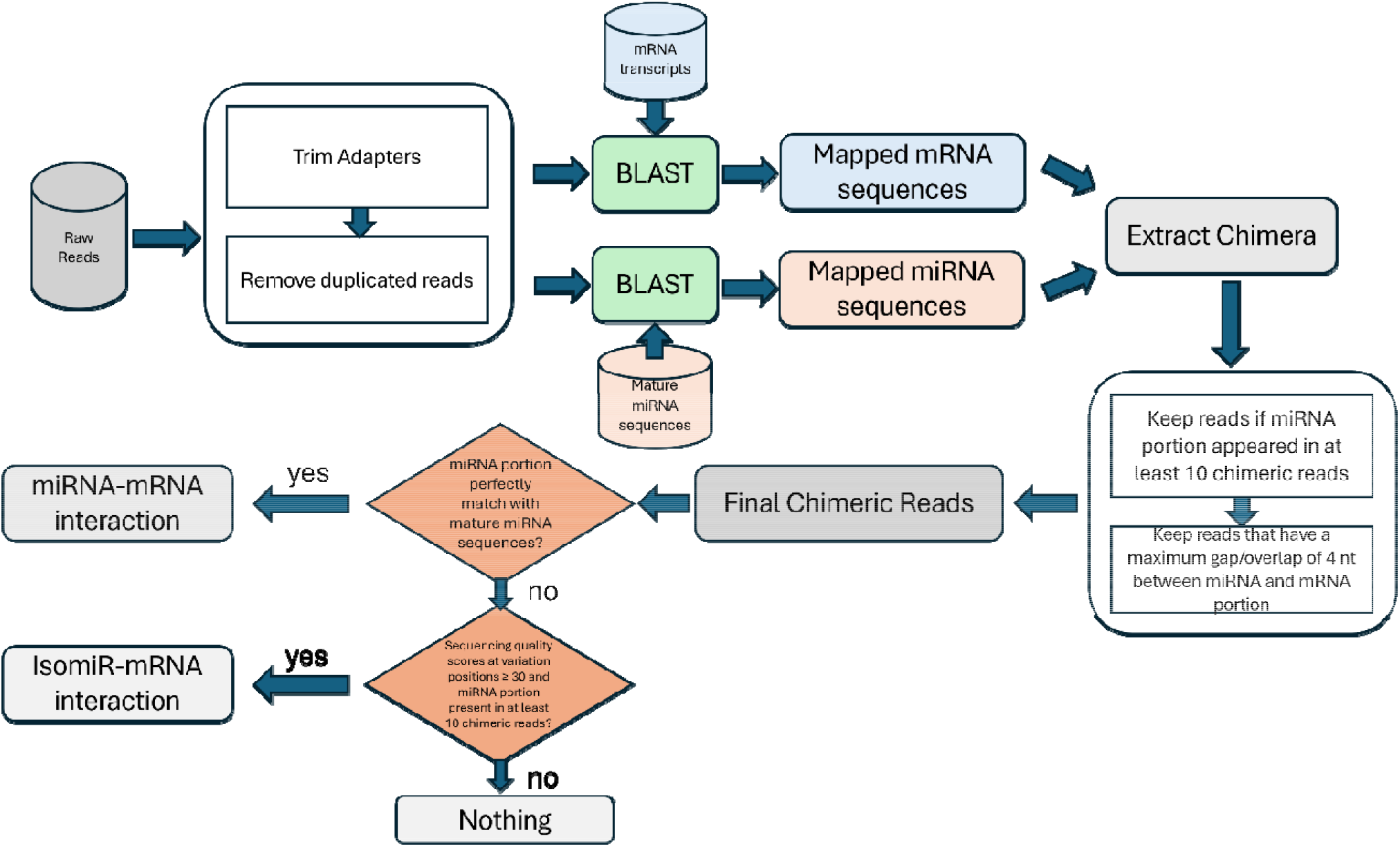
The pipeline for extracting miRNA/isomiR-mRNA interactions.

Each read mapped to both mRNA and miRNA was considered as a chimeric read. To ensure the quality of the miRNA portion in a chimeric read, which was typically shorter and more prone to sequencing errors, we retained only those chimeric reads that had the same miRNA segments as at least nine other chimeric reads. In addition, we excluded chimeric reads with more than a four-nucleotides gap or overlap between the mRNA and miRNA portions. Given that both miRNA and mRNA portions of a chimeric read could map to multiple miRNA and mRNA transcripts, we applied the following criteria sequentially to retain the most significant miRNA–mRNA pair: (1) the pair with the lowest BLAST E-values, and (2) if e-values were identical, the pair with the highest BLAST bit score. In total, we identified 5,556,801 chimeric reads across the 67 samples from the six datasets (Supplementary Table S1) (Fig. 1).

### Identify isomiRs and miRNA/isomiR-mRNA interactions

Following the procedure outlined above, each chimeric read consists of two segments: a mRNA portion segment and a miRNA portion segment, with a maximum overlap of four nucleotides between them. For each chimeric read, we classify the miRNA portion as an miRNA if it is an exact substring of a mature miRNA sequence in miRbase^25^. If it does not match a mature miRNA perfectly, we consider it an isomiR, provided this exact miRNA segment appears in at least ten chimeric reads (Fig. 1). This threshold of ten reads helps ensure that the likelihood of a BLAST hit being a random artifact is less than 8.9E-8, a cutoff also used in DMISO^15^. Unlike DMISO, however, when the miRNA segment only partially matches a mature miRNA, we extend the isomiR to the full-length miRNA sequence by appending the remaining unrepresented portion of the miRNA. This extension accounts for the uncertainty in whether the miRNA is truncated during biogenesis (such as in the case of a 3’ isomiR) or represents a full-length miRNA/isomiR, given the nature of the CLASH-like experiments. For the mRNA portion of each chimeric read, it must exactly match a string in the mRNA sequences annotated by GENCODE^24^. This approach allowed us to extract all miRNA-mRNA and isomiR-mRNA interactions from each sample in every dataset.

### The extracted isomiRs in isomiRdb

We downloaded the isomiR repertoire table from isomiRdb^9^. This table links each sequence to its corresponding mature miRNA name and assigns an isoLabel indicating whether the sequence is “canonical” or variants. We classified all sequences labeled as “canonical” in isomiRdb as miRNA sequences, while those with other isoLabels were considered isomiRs. This yielded a total of 2,573 annotated miRNAs and 3,274,688 annotated isomiRs. The number of isomiRs associated with each miRNA ranged from 5 to 45,751, with the median, mean, and standard deviation of isomiR counts being 258, 1,233, and 3,549, respectively. We then compared the extracted isomiRs from the six datasets with the annotated isomiRs in isomiRdb to assess whether a predicted isomiR was already annotated and explored the difference between the annotated isomiRs/miRNAs and the remaining unannotated ones.

### Extracted miRNA-mRNA interactions in miRTarBase

We downloaded the miRNA-mRNA interaction data in human from miRTarBase^20^ (hsa_MTI.csv file). The dataset contained 2,975 human miRNAs, which collectively targeted 16,976 mRNAs. It is important to note that some of these miRNAs, particularly from earlier annotation versions, may be inaccurate, as more recent annotations in miRBase include fewer miRNAs^25^. The number of mRNA targets per miRNA ranged from 1 to 3,238, with the median, mean, and standard deviation of mRNA targets being 354, 581, and 579, respectively. We then compared the extracted isomiR–mRNA and miRNA–mRNA interactions with those annotated in miRTarBase. For this comparison, we considered an isomiR–mRNA interaction as equivalent to a miRNA–mRNA interaction if the isomiR was a variant of the corresponding miRNA.

### Properties of extracted isomiRs/miRNAs and their interactions with mRNAs

We analyzed the nucleotide composition of miRNAs with a high number of extracted isomiRs compared with that of miRNAs with fewer extracted isomiRs. For each miRNA, we calculated the percentage of each nucleotide type. We then applied the Mann-Whitney test^27^ to compare the two groups of percentages to see whether the percentages in one group were different from those in the other group. Similarly, we compared the nucleotide content of miRNAs with a high number of extracted targets with that of miRNAs with fewer targets. We also examined the nucleotide content of common matching 6-mers between miRNAs that had matching seed regions and miRNAs that did not. A common matching 6-mer was defined as a six-nucleotide RNA segment that was present in both an isomiR/miRNA and more than 90% of its mRNA targets. We considered a miRNA/isomiR having a matching seed region if it had a common matching 6-mer started at positions 2 to 4 of this miRNA/isomiR sequence.

We further examined preferred mutation positions in miRNAs using a binomial test. A mutation position was defined as a nucleotide site within the mature miRNA where at least 5% of its associated isomiRs exhibited a variation. For each miRNA with *n* isomiRs, if a specific position was mutated in *k* of them, we computed a *p*-value as the binomial tail probability of observing *k* or more isomiRs with mutations at that position, under the null hypothesis that mutations occur uniformly at random across all 22 nucleotide positions of this mature miRNA. Positions with *p*-values less than 0.05 were considered statistically significant, indicating that the corresponding miRNA exhibited a preferred mutation site at the corresponding position.

## Results

### Identification of over four hundred novel isomiRs

We identified 1,233 miRNAs interacting with 17,845 mRNAs across six datasets (Table 1). Among these, 407 miRNAs had at least one corresponding isomiR. Notably, 100 of these 407 miRNAs were represented exclusively by isomiRs, with no reference miRNA sequences detected. In total, we identified 1,747 unique isomiRs that appeared at least 10 times across the datasets. The mean, median, and standard deviation of the number of chimeric reads supporting each isomiR were 251, 37, and 2,955, respectively. This distribution, with a low median but high standard deviation, indicated that while most isomiRs occurred only a few dozen times, a small subset appeared extremely frequently. Specifically, 1,298 isomiRs were supported by 10–99 reads, 395 by 100–999 reads, 49 by 1,000–9,999 reads, and five by more than 10,000 reads.

**Table 1.** The number of miRNAs, isomiRs, and their interacting mRNAs across datasets.

| Dataset | Samples | miRNAs | isomiRs | mRNAs | miRNA-mRNA interactions | isomiR-mRNA interactions |
| --- | --- | --- | --- | --- | --- | --- |
| CLASH | 9 | 405 | 111 | 9219 | 63126 | 15379 |
| CLEAR-CLIP | 12 | 262 | 152 | 2549 | 17617 | 5540 |
| qCLASH | 9 | 543 | 431 | 15549 | 2397074 | 104048 |
| eCLIP | 13 | 812 | 1466 | 17039 | 1776264 | 221029 |
| Kozer | 3 | 511 | 286 | 14305 | 601513 | 63280 |
| Fields | 21 | 525 | 173 | 11278 | 262192 | 29739 |
| Total | 67 | 1233 | 1747 | 17845 | 5117786 | 439015 |

We compared our identified isomiRs to those curated in isomiRdb^9^. While isomiRdb included 3,274,688 isomiRs from 2,573 miRNAs, our study identified 1,747 isomiRs from 1,233 miRNAs. Of these, 1,277 (73.1%) were also found in isomiRdb, suggesting that the extracted isomiRs are highly reliable. Conversely, only 0.03% of isomiRs in isomiRdb were recovered in our study, reflecting the limited experimental scope and narrower biological conditions reflected by the six datasets. To enable a fairer comparison, we focused on the 1,197 miRNAs common to both datasets. Within this subset, our 1,747 isomiRs were compared to 2,604,710 isomiRs in isomiRdb. Again, 1,277 (73.1%) of our isomiRs matched entries in isomiRdb, while representing only 0.05% of the isomiRdb set.

The remaining 26.9% (470) of our identified isomiRs not found in isomiRdb are still likely to be biologically valid. This confidence is supported by two key observations. First, these isomiRs were observed repeatedly across samples and datasets. On average, each unsupported isomiR appeared in 4.0 distinct samples and 1.4 dataset, compared with 3.4 samples and 1.3 datasets for the supported isomiRs. Second, the read counts supporting these 26.9% of isomiRs were similar to those supporting isomiRs already present in isomiRdb. The median and mean number of chimeric reads for supported isomiRs were 38 and 254, respectively, compared to 33 and 245 for unsupported ones. These similarities in the number of samples, datasets and chimeric reads strongly suggest that the unsupported isomiRs are equally reliable and that existing isomiR annotations remain incomplete.

To further investigate the highly abundant isomiRs, we examined the 54 isomiRs supported by more than 1,000 chimeric reads. To rule out experimental artifacts, we assessed their distribution across datasets. These isomiRs were present in an average of 23 distinct samples from multiple datasets, suggesting that their abundance reflects true biological activity across experimental conditions rather than noise or artifacts of individual datasets and samples. We also examined whether the reference miRNAs of these 54 isomiRs had similar chimeric read abundance. These isomiRs originated from 33 reference miRNAs. Interestingly, 28 of these miRNAs also produced at least one other isomiR supported by fewer than 100 reads, suggesting differential isomiR expression from the same miRNA precursor. Of the remaining five miRNAs, four had only one detected isomiR, while the fifth, hsa-miR-30b-3p, had a second isomiR supported by over 200 reads. Notably, these 33 miRNAs had a median of 13 and a mean of 18.7 associated isomiRs, underscoring the diversity and differential expression of isomiRs derived from a single miRNA.

### miRNAs with higher adenine and thymine but lower cytosine content tend to have more isomiRs and mRNA targets

As noted above, we identified at least one isomiR for each of the 407 miRNAs, with 100 of them lacking the reference miRNA detection and represented solely by isomiRs. The number of isomiRs per miRNA ranged from 1 to 101, with a mean of 4 and a median of 1. Specifically, 227, 53, 22, 14, and 91 miRNAs were associated with one, two, three, four, and at least five isomiRs, respectively.

To explore why certain miRNAs give rise to more isomiRs, we examined the nucleotide composition of their reference sequences (Materials and Methods). We found that miRNAs with more isomiRs tend to have higher adenine and thymine content and lower cytosine content (Supplementary Table S2). For example, when comparing miRNAs with one to four isomiRs to those with five or more, the differences in adenine, thymine, and cytosine content were statistically significant, with p-values of 0.015, 0.009, and 0.010, respectively (Table 2).

**Table 2.**
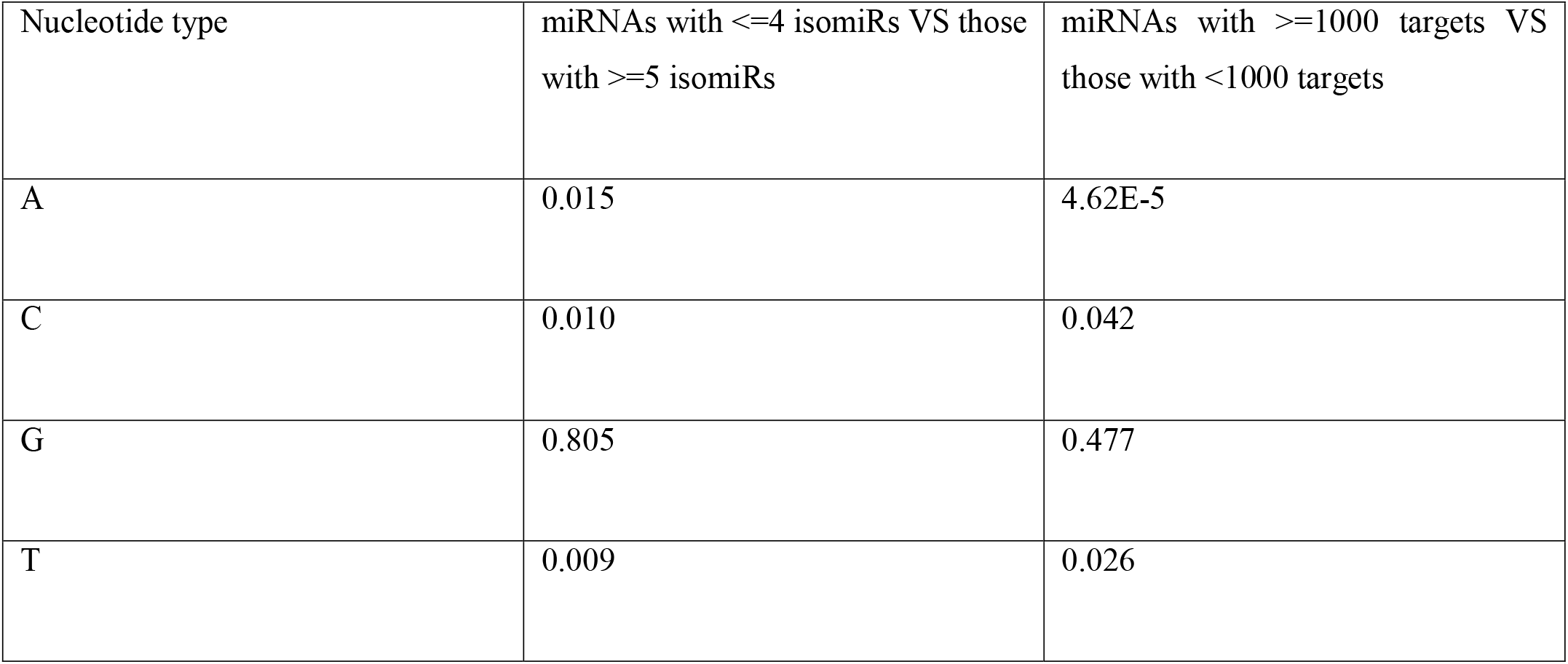
Nucleotide preference of miRNAs with more isomiRs and more targets.

We also investigated whether certain nucleotide positions in miRNAs were more prone to mutations that give rise to isomiRs (Materials and Methods). Our analysis showed that the seed region was relatively conserved, with mutations occurring less frequently than in non-seed regions. In contrast, mutation rates were significantly higher outside the seed region, with 12 non-seed positions showing a p-value < 0.05 and 10 positions with p-value < 0.01. The mutation frequency exhibited a bimodal distribution, with notable peaks at positions 10 and 16 (Supplementary Table S3 and Figure S1).

Considering miRNAs and their associated isomiRs together, the number of mRNA targets per miRNA varied widely, from 1 to 12,910, with a mean of 697 and a median of 28. The high standard deviation (1,843) reflects the substantial variation in target counts among different miRNAs. In detail, 810 miRNAs had between 1 and 99 targets, 250 had 100 to 999 targets, 164 had 1,000 to 9,999 targets, and 9 miRNAs had over 10,000 targets.

To understand what distinguished miRNAs with many targets from those with fewer, we compared the nucleotide composition of 173 miRNAs with ≥1,000 targets to that of the remaining 1,060 miRNAs with fewer targets, using the Mann-Whitney test^27^. We found that miRNAs with many targets had significantly more adenines (p < 4.62×10□□) and thymines (p < 0.05), and significantly fewer cytosines (p < 0.05), again supporting the nucleotide bias observed for isomiR-rich miRNAs (Table 2).

We next focused on the nine miRNAs with more than 10,000 mRNA targets. All but one of these miRNAs (hsa-miR-3168) belonged to miRNA families with at least two homologous members, and six had three or more homologs in miRBase^25^ (Supplementary Table S4). We defined homologous miRNAs as those with identical mature sequences located in different genomic regions, such as hsa-let-7a-5p and hsa-let-7f-5p. Given that only 268 of the 1,233 miRNAs in our study have three or more homologs in miRBase, the probability of observing six of the nine high-target miRNAs with such homology is statistically significant (p = 0.005). This suggests that miRNA family expansion contributes to broader target coverage.

We also assessed whether these nine miRNAs with most targets had more isomiRs than others. Remarkably, five of them had between 30 and 60 isomiRs each. Since only seven of the 1,233 miRNAs in our dataset had 30 or more isomiRs, the probability of five of them appearing among the nine is extremely low (p = 1.12×10□^1^□). This result reinforces the idea that miRNA family size contributes to both isomiR diversity and target multiplicity. Although hsa-miR-3168 lacked homologs, it exhibited other characteristics that might explain its large target set: it was shorter (17 nucleotides compared to the typical 22) and had a higher adenine content (0.29 vs. an average of 0.22). These features may enhance its ability to bind a wider range of mRNA targets.

### IsomiRs significantly expand the target repertoire of their reference miRNAs

Chimeric reads provide direct evidence of interactions between miRNAs/isomiRs and their target mRNAs. Across the six datasets analyzed, we identified 830,336 unique miRNA–mRNA interactions and 132,530 unique isomiR–mRNA interactions. To assess the reliability and novelty of these interactions, we compared our findings with the curated interactions in miRTarBase (Materials and Methods).

In total, our study identified 1,233 miRNAs targeting 17,845 mRNAs. Of these, 1,184 miRNAs and 9,451 mRNAs were also present in miRTarBase. These overlapping entities were involved in 1,730,720 curated interactions in miRTarBase and 858,788 interactions in our dataset. We found that 57,861 of the curated interactions were also detected in our study (recall = 0.067), while 57,861 of our detected interactions were present in miRTarBase (precision = 0.033). These relatively low recall and precision values may reflect several factors: the incompleteness of current miRTarBase annotations, the mis-annotations in current miRTarBase (e.g., the 2,975 miRNAs used), the limited sample diversity in our study, and the condition-specific nature of miRNA/isomiR–mRNA interactions. Interestingly, the interactions found in both our study and miRTarBase occurred in an average of 4.2 samples, suggesting that these interactions are reproducible and potentially biologically relevant. Notably, 14.8% of the interactions not supported by miRTarBase were observed in more than 4.2 samples, implying that many of these unsupported interactions may be genuine but uncurated.

We next investigated the extent to which isomiRs contribute independently to miRNA–mRNA targeting. We focused on the 307 miRNAs for which both the reference miRNA and at least one isomiR were present in our datasets. We found that, on average, 18.9% of the targets for each miRNA–isomiR group were uniquely attributable to isomiRs, with a median of 2.7%. For example, hsa-miR-30b-3p and its two isomiRs jointly targeted 629 mRNAs. Of these, 46 (7.3%) were unique to the reference miRNA, 539 (85.7%) were unique to the isomiRs, and only 44 (7.0%) were shared. Strikingly, in 47 (15.3%) of the 307 cases, isomiRs contributed more unique targets than the corresponding reference miRNAs. We noticed that these miRNAs, target mRNAs, and isomiRs did occur in samples, suggesting that the absence of reference miRNA binding in some cases was not due to the lack of expression of the miRNA or the mRNA. This points to a biologically meaningful shift in targeting specificity, emphasizing the need to account for isomiRs in target prediction.

To explore why certain targets were bound exclusively by isomiRs, we analyzed seed region sequences. Among the 307 miRNAs, 301 had isomiRs that bound targets not bound by the reference miRNA. These 301 miRNAs produced 1,265 distinct isomiRs with unique targets. We found that 304 (24.0%) of these isomiRs, corresponding to 149 (49.5%) of the 301 miRNAs, had at least one nucleotide alteration in their seed regions. Surprisingly, the remaining 961 isomiRs (76.0%), associated with 152 miRNAs (50.5%), had unchanged seed sequences compared to their reference miRNAs. This suggests that in many cases, regions outside the seed may play a dominant role in isomiR-mediated targeting, as further explored in the next section.

### A subset of miRNAs and isomiRs prefer non-seed binding to their targets

Previous studies have shown that certain miRNAs can bind their targets through regions outside the seed region (positions 2–7), yet the prevalence and characteristics of such non-seed binding remain unclear^9,12-14,28-30^. The collected chimeric reads provided a unique opportunity to systematically investigate the extent of non-seed binding among miRNAs and isomiRs. To address this, we analyzed 1,133 miRNAs and 1,747 isomiRs identified across the six datasets, excluding the 100 miRNAs for which only isomiRs were detected. We focused on a subset of 862 miRNAs and 1,500 isomiRs, each with at least six mRNA targets. This cutoff ensured that the false discovery rate was controlled at below one false positive per miRNA/isomiR.

For each miRNA or isomiR, we performed local alignments between all possible 6-mers within the miRNA/isomiR sequence and the reverse complement of its corresponding mRNA targets. For a miRNA or isomiR, we retained only those 6-mers that matched more than 90% of its associated target sequences. These frequently observed 6-mers were considered the common binding motifs of a given miRNA or isomiR^31^. Using this approach, we found that 544 miRNAs and 1,405 isomiRs had at least one such conserved 6-mer motif. Among these, 435 miRNAs (80.0%) and 1,290 isomiRs (91.8%) had their common 6-mers overlapping with the canonical seed region (defined as sharing at least three nucleotides with positions 2–7 of the mature miRNA sequence). Consequently, 20.0% of miRNAs and 8.2% of isomiRs appear to bind their targets without engaging the seed region. Furthermore, we found that seed-binding miRNAs and isomiRs frequently used additional non-seed regions for targeting. Specifically, 94.5% of seed-binding miRNAs and 95.7% of seed-binding isomiRs also had at least one non-seed 6-mer motif, suggesting that seed-based recognition is often complemented by non-seed interactions.

We then compared the characteristics of seed-binding and non-seed-binding miRNAs/isomiRs in four ways: 1). Number of matching 6-mers: Seed-binding miRNAs had significantly more matching 6-mers (mean and median = 9) compared to non-seed-binding miRNAs (mean = 3, median = 2). The difference was highly significant (Mann–Whitney p = 1.0E–37). Location of matching regions: In non-seed-binding miRNAs/isomiRs, we observed that over half of the matching regions began between positions 6 and 8. Additionally, position 11 was part of the matching region in 73.5% of these miRNAs/isomiRs, highlighting the importance of the central region in non-seed-mediated targeting; 3). Overall nucleotide composition: We found no significant difference in the overall nucleotide composition between seed-binding and non-seed-binding miRNAs/isomiRs; 4). Nucleotide composition of matching regions: When analyzing the nucleotide content within the common 6-mers, we found that seed-binding motifs were significantly enriched in adenine and thymine and depleted in cytosine compared to non-seed-binding motifs (Mann–Whitney p < 0.05). This finding is consistent with earlier observations linking higher adenine/thymine content to increased numbers of isomiRs and mRNA targets.

Together, these results indicate that while seed-based recognition is predominant, a substantial fraction of miRNAs and isomiRs utilize non-seed regions, particularly their central region, for target binding. The characteristics of these non-canonical interactions suggest potential mechanisms of regulatory flexibility and underscore the importance of considering full-length miRNA and isomiR sequences in target prediction models^29^.

## Discussion

In this study, we analyzed 67 human CLASH and CLASH-like samples and identified 1,747 unique isomiRs and 439,015 isomiR–mRNA interactions. Approximately three-quarters of these isomiRs were previously annotated in isomiRdb^9^, while only about 6.7% of the identified miRNA/isomiR–mRNA interactions were found in miRTarBase^20^. We found that miRNAs enriched in adenine and thymine but depleted in cytosine tended to generate more isomiRs and had a greater number of mRNA targets. On average, 19.0%, and in some cases, up to 98.1%, of a miRNA’s targets were attributable exclusively to its isomiRs, even though many of these isomiRs shared identical seed sequences with their corresponding reference miRNAs.

Our results highlight significant gaps in the current understanding of isomiRs and their regulatory roles. Despite analyzing only 67 samples, 26.9% of the extracted isomiRs were not present in isomiRdb. These unannotated isomiRs occurred with similar frequency and across a comparable number of samples as those previously cataloged in isomiRdb, suggesting that existing isomiR annotations remain incomplete. Likewise, only a small fraction (6.7%) of the detected miRNA/isomiR–mRNA interactions overlapped with those curated in miRTarBase, and curated interactions were not better supported by the number of chimeric reads and samples than non-curated ones. Together, these findings suggest that the true diversity and impact of isomiR-mediated regulation are likely underestimated.

Although we attempted to include all available CLASH and CLASH-like datasets, the analysis was limited to 67 samples. As more such datasets become available in the future, the patterns and conclusions drawn here can be revisited and validated on a larger scale. Moreover, current experimental technologies often fail to capture the full-length sequences of miRNAs/isomiRs and their complete target mRNA fragments. Improved methodologies that preserve these full-length chimeric reads will provide more precise insights into the molecular mechanisms of miRNA/isomiR targeting. Finally, due to the lack of functional assays or complementary datasets, this study could not assess the downstream biological effects of the identified interactions. Future studies incorporating transcriptomic, proteomic, or reporter assay data may help to better elucidate the functional significance of miRNA and isomiR-mediated regulation.

## Acknowledgements

This work has been supported by the National Science Foundation [Grants 2120907 and 2015838].

## Author Contributions

H.H. and X.L. conceived and designed the study. M.W. performed the experiments. M.W., R.C.R., X.L., and H.H. analyzed the data. M.W., R.C.R., X.L., and H.H. wrote the manuscript. All authors proofread the manuscript.

## Data availability

The CLASH, CLEAR-CLIP, qCLASH, chimeric eCLIP, and Fields et al. data were downloaded, respectively, from accession numbers GSE50452, GSE73059, GSE101978, GSE198251, and GSE164634. The Kozer et al. data is downloaded from https://www.ebi.ac.uk/biostudies/arrayexpress/studies/E-MTAB-8591. The Supplementary Table S1 is at https://figshare.com/articles/dataset/Table_S1/28958909?file=54310421. The Supplementary Tables S2 - S4 are at https://figshare.com/articles/dataset/Supplementary_tables_S2_-_S4/28959281. The Supplementary Figure S1 is available at https://figshare.com/articles/figure/Figure_S1/28959335?file=54312092.

## References

1 Morin, R. D. et al. Application of massively parallel sequencing to microRNA profiling and discovery in human embryonic stem cells. Genome research 18, 610–621, doi:10.1101/gr.7179508 (2008).

2 Cloonan, N. et al. MicroRNAs and their isomiRs function cooperatively to target common biological pathways. Genome biology 12, doi:10.1186/gb-2011-12-12-r126 (2011).

3 van der Kwast, R. V. C. T., Woudenberg, T., Quax, P. H. A. & Nossent, A. Y. MicroRNA-411 and Its 51-IsomiR Have Distinct Targets and Functions and Are Differentially Regulated in the Vasculature under Ischemia. Mol Ther 28, 157–170, doi:10.1016/j.ymthe.2019.10.002 (2020).

4 Mercey, O. et al. Characterizing isomiR variants within the microRNA-34/449 family. Febs Lett 591, 693–705, doi:10.1002/1873-3468.12595 (2017).

5 Zelli, V. et al. Emerging Role of isomiRs in Cancer: State of the Art and Recent Advances. Genes-Basel 12, doi:10.3390/genes12091447 (2021).

6 van der Kwast, R. V. C. T., Quax, P. H. A. & Nossent, A. Y. An Emerging Role for isomiRs and the microRNA Epitranscriptome in Neovascularization. Cells-Basel 9, doi:10.3390/cells9010061 (2020).

7 Wagner, V., Meese, E. & Keller, A. The intricacies of isomiRs: from classification to clinical relevance. Trends in Genetics 40, 784–796, doi:10.1016/j.tig.2024.05.007 (2024).

8 Wang, Y., Goodison, S., Li, X. M. & Hu, H. Y. Prognostic cancer gene signatures share common regulatory motifs. Scientific reports 7, doi:ARTN 4750 10.1038/s41598-017-05035-3 (2017).

9 Aparicio-Puerta, E. et al. isomiRdb: microRNA expression at isoform resolution. Nucleic acids research 51, D179–D185, doi:10.1093/nar/gkac884 (2023).

10 Bofill-De Ros, X. et al. Tumor IsomiR Encyclopedia (TIE): a pan-cancer database of miRNA isoforms. Bioinformatics 37, 3023–3025, doi:10.1093/bioinformatics/btab172 (2021).

11 Zhang, Y. W. et al. IsomiR Bank: a research resource for tracking IsomiRs. Bioinformatics 32, 2069–2071, doi:10.1093/bioinformatics/btw070 (2016).

12 Nersisyan, S. et al. isomiRTar: a comprehensive portal of pan-cancer 5’-isomiR targeting. Peerj 10, doi:10.7717/peerj.14205 (2022).

13 Chen, Y. H. & Wang, X. W. miRDB: an online database for prediction of functional microRNA targets. Nucleic acids research 48, D127–D131, doi:10.1093/nar/gkz757 (2020).

14 McGeary, S. E. et al. The biochemical basis of microRNA targeting efficacy. Science 366, 1470–+, doi:10.1126/science.aav1741 (2019).

15 Talukder, A., Zhang, W. C., Li, X. M. & Hu, H. Y. A deep learning method for miRNA/isomiR target detection. Scientific reports 12, doi:10.1038/s41598-022-14890-8 (2022).

16 Helwak, A., Kudla, G., Dudnakova, T. & Tollervey, D. Mapping the Human miRNA Interactome by CLASH Reveals Frequent Noncanonical Binding. Cell 153, 654–665, doi:10.1016/j.cell.2013.03.043 (2013).

17 Gay, L. A., Turner, P. C. & Renne, R. Modified Cross-Linking, Ligation, and Sequencing of Hybrids (qCLASH) to Identify MicroRNA Targets. Curr Protoc 1, doi:10.1002/cpz1.257 (2021).

18 Manakov, S. et al. Scalable and deep profiling of mRNA targets for individual microRNAs with chimeric eCLIP. (2022).

19 Moore, M. J. et al. miRNA-target chimeras reveal miRNA 31-end pairing as a major determinant of Argonaute target specificity. Nature communications 6, doi:10.1038/ncomms9864 (2015).

20 Cui, S. D. et al. miRTarBase 2025: updates to the collection of experimentally validated microRNA-target interactions. Nucleic acids research 53, D147–D156, doi:10.1093/nar/gkae1072 (2024).

21 Kozar, I. et al. Cross-Linking Ligation and Sequencing of Hybrids (qCLASH) Reveals an Unpredicted miRNA Targetome in Melanoma Cells. Cancers 13, doi:10.3390/cancers13051096 (2021).

22 Fields, C. J. et al. Sequencing of Argonaute-bound microRNA/mRNA hybrids reveals regulation of the unfolded protein response by microRNA-320a. PLoS genetics 17, doi:10.1371/journal.pgen.1009934 (2021).

23 Bolger, A. M., Lohse, M. & Usadel, B. Trimmomatic: a flexible trimmer for Illumina sequence data. Bioinformatics 30, 2114–2120, doi:10.1093/bioinformatics/btu170 (2014).

24 Harrow, J. et al. GENCODE: the reference human genome annotation for The ENCODE Project. Genome research 22, 1760–1774, doi:10.1101/gr.135350.111 (2012).

25 Kozomara, A., Birgaoanu, M. & Griffiths-Jones, S. miRBase: from microRNA sequences to function. Nucleic acids research 47, D155–D162, doi:10.1093/nar/gky1141 (2019).

26 Altschul, S. F., Gish, W., Miller, W., Myers, E. W. & Lipman, D. J. Basic Local Alignment Search Tool. Journal of molecular biology 215, 403–410, doi:10.1016/S0022-2836(05)80360-2 (1990).

27 Mann, H. B. & Whitney, D. R. On a Test of Whether one of Two Random Variables is Stochastically Larger than the Other. Annals of Mathematical Statistics 18, 50–60, doi:10.1214/aoms/1177730491 (1947).

28 Ding, J., Li, X. M. & Hu, H. Y. CCmiR: a computational approach for competitive and cooperative microRNA binding prediction. Bioinformatics 34, 198–206, doi:10.1093/bioinformatics/btx606 (2018).

29 Talukder, A., Li, X. M. & Hu, H. Y. Position-wise binding preference is important for miRNA target site prediction. Bioinformatics 36, 3680–3686, doi:10.1093/bioinformatics/btaa195 (2020).

30 Athaya, T., Li, X. M. & Hu, H. Y. A deep learning method to integrate extracelluar miRNA with mRNA for cancer studies. Bioinformatics 40, doi:10.1093/bioinformatics/btae653 (2024).

31 Ding, J., Cai, X., Wang, Y., Hu, H. & Li, X. ChIPModule: systematic discovery of transcription factors and their cofactors from ChIP-seq data. Pacific Symposium on Biocomputing. Pacific Symposium on Biocomputing, 320–331 (2013).

